# *PySTARC*: GPU-accelerated Brownian dynamics for bimolecular association rate constants

**DOI:** 10.64898/2026.09.28.754770

**Authors:** Anupam Anand Ojha, Gary Huber, Shiksha Dutta, Sonya M. Hanson

## Abstract

Drug-target association and dissociation rates often determine *in vivo* efficacy more than affinity alone. The association rate *k*_on_, however, is computationally challenging to predict since the productive encounter is a rare event in a large translational and orientational search space, which lies beyond the reach of conventional atomistic simulations. We present *PySTARC* (*Py*thon *S*imulation *T*oolkit for *A*ssociation *R*ate *C*onstants), a GPU-accelerated Brownian dynamics engine for estimating bimolecular association rate constants. *PySTARC* is a Python reimplementation of the BrownDye engine that converges even small reaction probabilities on a single GPU, at a throughput that would otherwise require a large CPU cluster. The current framework resolves reactions between integration steps with a closed-form Brownian bridge, models the internal flexibility of the solute through a coarse-grained bead chain, distributes trajectories across multiple GPUs, checkpoints long runs, monitors convergence, and automates the entire workflow from input structures to rate estimates. *PySTARC* is validated against protein-ligand and protein-protein complexes spanning five orders of magnitude in *k*_on_, with most estimates within one order of magnitude of experiment.

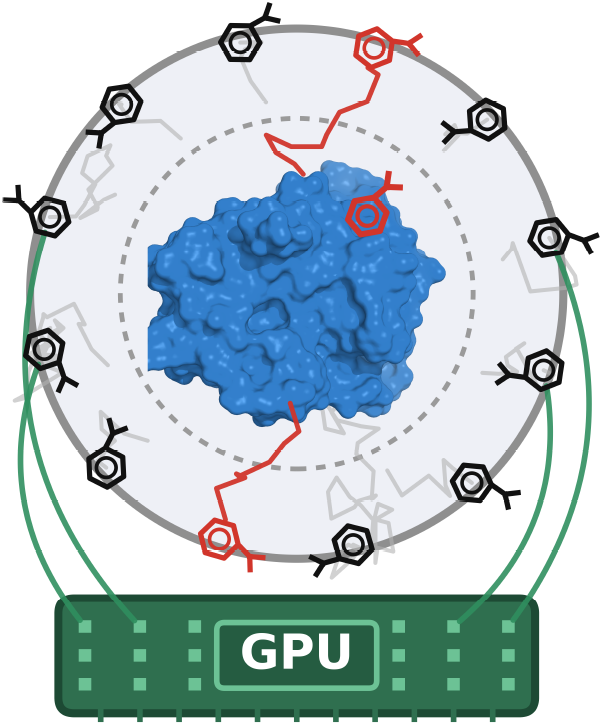

## INTRODUCTION

Drug-target affinity is measured at equilibrium as the dissociation constant *K*_*d*_, i.e., the ratio of the dissociation (*k*_off_) and association (*k*_on_) rate constants. Ligands with the same *K*_*d*_ can therefore have *k*_on_ and *k*_off_ that differ by orders of magnitude ^1^. Despite this degeneracy, lead optimization has long relied on *K*_*d*_ and the binding free energy Δ*G*_bind_ to guide structure-activity relationships ^1–6^. Physics-based free energy methods span alchemical relative binding free energies (FEP+) ^7,8^, thermodynamic integration ^9,10^, endpoint approximations such as MM/PBSA and MM/GBSA ^11,12^, and absolute binding free energy calculations by double decoupling and the alchemical transfer method ^13,14^, while deep learning affinity predictors, from convolutional neural networks such as K_DEEP_ ^15^, Pafnucy ^16^, OnionNet ^17^, and GNINA ^18^ to graph neural networks such as PIGNet ^19^ and GIGN ^20^, and most recently, the co-folding model Boltz-2 that predicts both the complex structure and its affinity ^21^, estimate binding affinity directly from structure, without the explicit conformational sampling required by rigorous physics-based methods. Kinetic characterization, by contrast, quantifies the rate of complex formation through *k*_on_ and the complex life-time through the residence time *τ* = 1 / *k*_off_, and can be more predictive of *in vivo* efficacy than *K*_*d*_ alone ^2,6,22^. Methodological development has largely targeted the estimation of *k*_off_ and the residence time ^5,23,24^, through enhanced sampling frameworks such as metadynamics and its infrequent sampling variant ^25,26^, random acceleration molecular dynamics (RAMD) ^27,28^, targeted molecular dynamics ^29,30^, Markov state models ^31–33^, weighted ensemble path sampling in both its original and later resampling formulations, ^34–38^ and milestoning simulations with their quantum mechanical and machine learning augmented extensions ^39–43^.

The association rate constant *k*_on_ is bounded by the Smoluchowski diffusion limit 4*πN*_A_*DR* ≈ 10^9^ M^−1^ s^−1^ for a relative diffusion coefficient *D* and contact distance *R* ^45–47^. A drug that associates at the Smoluchowski limit gains affinity only by dissociating more slowly ^1,2^. Affinity differences between ligands of the same target have therefore been attributed largely to *k*_off_ ^22,48^. However, experimental association rate constants of different ligands span 10^3^-10^9^ M^−1^ s^−1 49^. Differences in *k*_on_ therefore contribute to affinity differences alongside *k*_off_. Association requires the drug and its target to diffuse together into an encounter complex, from which they either form the bound complex or separate and diffuse back into bulk solvent. As a consequence, *k*_on_ reflects both the rate of encounter and the probability of capture ^50–53^. Drug-target association is intrinsically more demanding to sample with atomistic molecular dynamics (MD) simulations than unbinding, since productive encounters occur within a large translational and orientational search space with no predefined reaction coordinate. In contrast, unbinding events are seeded from the known bound pose along a comparatively constrained exit path. A direct reconstruction of a binding event by unbiased atomistic simulations therefore requires either large ensembles of aggregated trajectories ^54^ or long timescale simulations on specialized hardware ^55^. Brownian dynamics (BD) simulations resolve *k*_on_ at a lower cost by treating the two solutes as rigid bodies whose relative diffusion is propagated subject to the electrostatic, desolvation, and hydrodynamic interactions between them ^44,56–60^. In BD, *k*_on_ is the product of the diffusion-controlled encounter rate and the reaction probability, which is the fraction of simulated encounters that satisfy the reaction criterion.

Established packages for estimating association rate constants, including but not limited to BrownDye ^58,62^, SDA ^59,63^, and GeomBD3^60^, remain confined to central processing units (CPUs) such that a buried or otherwise selective site, for which the reaction probability is small, requires a very large number of trajectories to converge and is therefore slow to compute ^64^. A comparable throughput limit once constrained all-atom MD simulations, which was overcome by moving the calculation onto graphics processing units (GPUs). Existing MD engines, such as OpenMM ^65^, AMBER ^66^, and GROMACS ^67,68^, brought the microsecond trajectory from a supercomputer to a single workstation, since the N-body force evaluation at each step maps onto the data-parallel architecture of the GPU ^69^. Unlike the coupled N-body force evaluation in MD simulations, the propagation of BD trajectories is mutually independent, an embarrassingly parallel structure that maps directly onto a single concurrent GPU batch.

We introduce *PySTARC* (*Py*thon *S*imulation *T*oolkit for *A*ssociation *R*ate *C*onstants), a GPU-accelerated BD engine that computes bimolecular association rate constants at high throughput. It is a Python reimplementation of the BrownDye engine ^58^, originally written in C++ and OCaml. BrownDye distributes its trajectories across CPU threads, while *PySTARC* advances all trajectories on GPUs such that even a small reaction probability converges at a fraction of the CPU cost. *PySTARC* adds several capabilities to the BD workflow. A closed-form Brownian-bridge crossing probability recovers reactions occurring between recorded steps. The reaction criterion is configurable and supports multi-state reaction schemes. The hydrodynamic, desolvation, and steric terms can each be enabled or disabled through a single configuration file. Simulation outputs are saved for downstream analysis, including trajectories, encounters, first-passage times, committors, contact and pose statistics, transition matrices, and convergence estimates. Long runs are checkpointed for restart, and an automated setup builds the simulation inputs directly from the complex structure. The same throughput makes *PySTARC* a source of consistent *k*_on_ labels for machine learning (ML) models of binding kinetics, which are limited by the scarcity of kinetic measurements relative to the affinity data behind modern scoring functions ^70–75^. The position and orientation of the diffusing solute at the binding criterion define the first-hitting point distribution ^76,77^, from which the encounter poses are reconstructed as a structural ensemble that seeds atomistic simulations where the rigid-body approximation no longer holds ^39^. The framework is validated against experimental rates for protein-ligand and protein-protein complexes whose *k*_on_ spans five orders of magnitude.

## THEORY AND IMPLEMENTATION

BD simulations estimate *k*_on_ within the Northrup-Allison-McCammon (NAM) framework ^44^, which partitions the region surrounding the receptor at the *b*-surface of radius *b* into an inner domain propagated explicitly and an outer domain treated analytically (Figure 1). The NAM algorithm factorizes the association rate constant into a diffusion-controlled encounter rate *k*_*b*_, at which the receptor and ligand reach the *b*-surface, and the probability *P*_rxn_ that an encounter proceeds to reaction,

**Figure 1:**
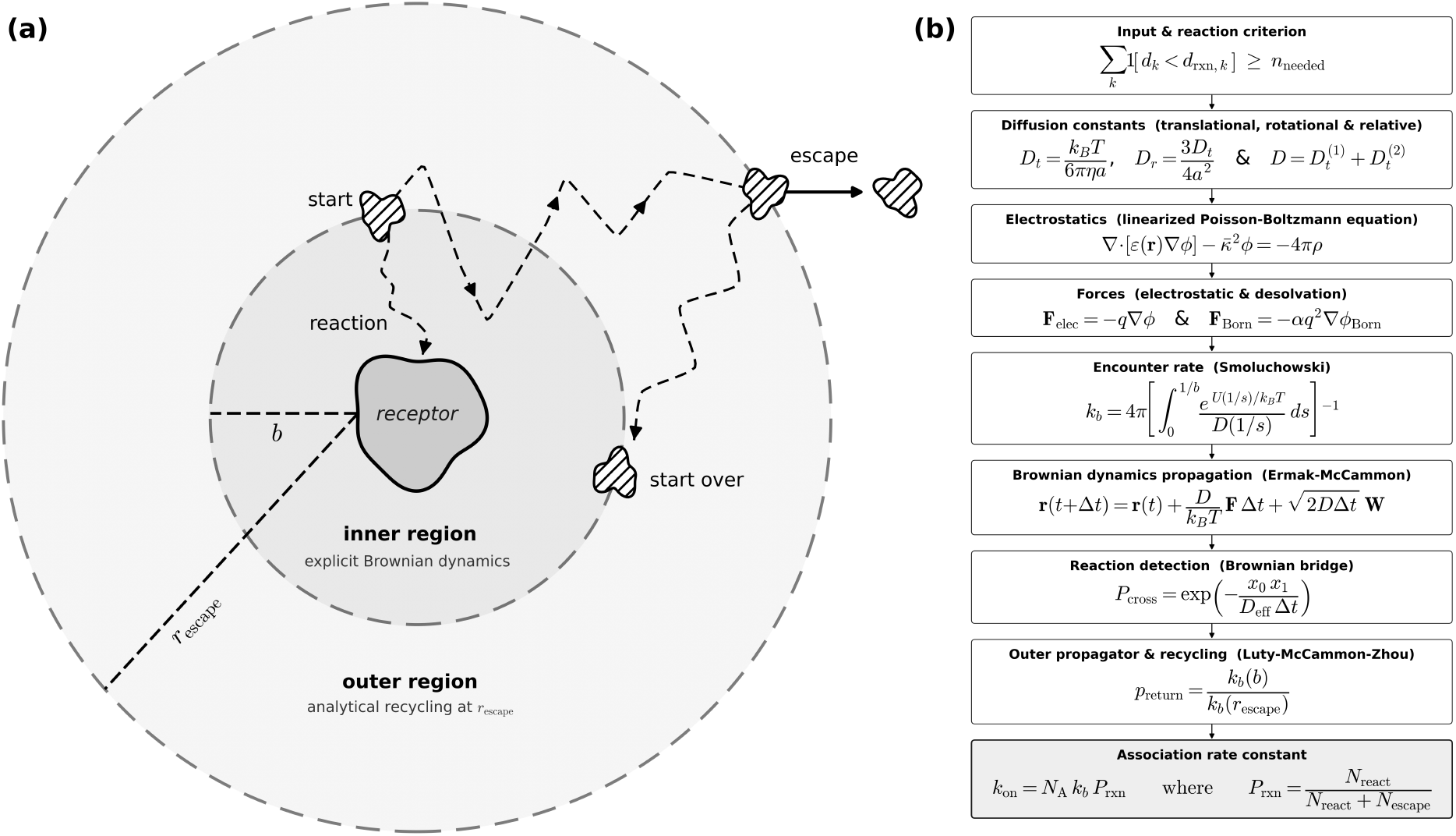
(a) In the Northrup-Allison-McCammon decomposition ^44^, the ligand is launched from the *b*-surface, a sphere of radius *b* enclosing the receptor. The ligand is then propagated relative to the fixed receptor by explicit Brownian dynamics until it either satisfies the reaction criterion at the receptor (*reaction*) or reaches the escape surface of radius *r*_escape_. At the escape surface, a trajectory is either terminated (*escape*) or returned to the *b*-surface (*start over*) with an analytically evaluated probability. *k*_on_ is obtained from the encounter rate *k*_*b*_ and the fraction of trajectories that react before escaping. Dashed curves are representative Brownian trajectories and hatched shapes denote the ligand. (b) Workflow for estimating *k*_on_ in *PySTARC*, with the governing equation at each step. A reaction is recorded when at least *n*_needed_ of the contact distances *d*_*k*_ fall below their cutoffs *d*_rxn,*k*_.

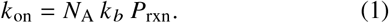

The encounter rate is computed from the potential of mean force *U*(*r*) and the relative diffusion coefficient *D*(*r*),

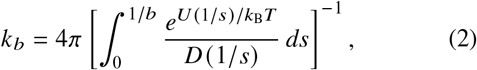

which reduces to 4*πDb* for two point charges (Supplementary Table S1). The reaction probability is estimated by launching independent trajectories on the *b*-surface and advancing each ligand under the electrostatic, desolvation, and hydrodynamic forces by the over-damped Ermak-McCammon step,

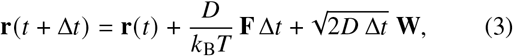

where **W** is a Gaussian noise vector of zero mean and unit variance. The forces are read from the linearized Poisson-Boltzmann and Born desolvation grids, the diffusion coefficients are obtained from the Stokes-Einstein relations with a Rotne-Prager-Yamakawa (RPY) correction at short range, and the timestep is adapted to the local gradient (Supplementary Table S1). The orientation is propagated by an analogous rotational step and represented as a quaternion. A trajectory is counted as reacted once the required contact pairs fall within their cutoffs, with crossings between successive steps recovered by a closed-form Brownian-bridge probability. A trajectory that reaches the escape surface without reacting is recycled with an analytically evaluated return probability or terminated as an escape (Figure 1a). The reaction probability is the reacted fraction,

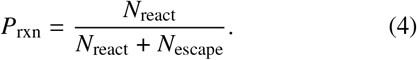

Trajectory propagation in *PySTARC* is carried out entirely on the GPUs (Figure 2). Independent of the trajectory count, the electrostatic and Born desolvation grids, the diffusion coefficients, and the reaction criterion are evaluated once on the CPUs and held on the GPUs for the duration of the run rather than recomputed per trajectory. For a rigid receptor, the force on the diffusing solute reduces to grid interpolation at every atomic site. A CUDA kernel launched via CuPy ^78^ performs the interpolation over a two-dimensional thread grid spanning trajectories and atomic sites. Across multiple GPUs, the ensemble is partitioned into disjoint shards, each propagated from an independently offset random seed. The reacted and escaped trajectory counts are returned to the CPUs and pooled, from which *k*_on_ and its confidence interval are computed.

**Figure 2:**
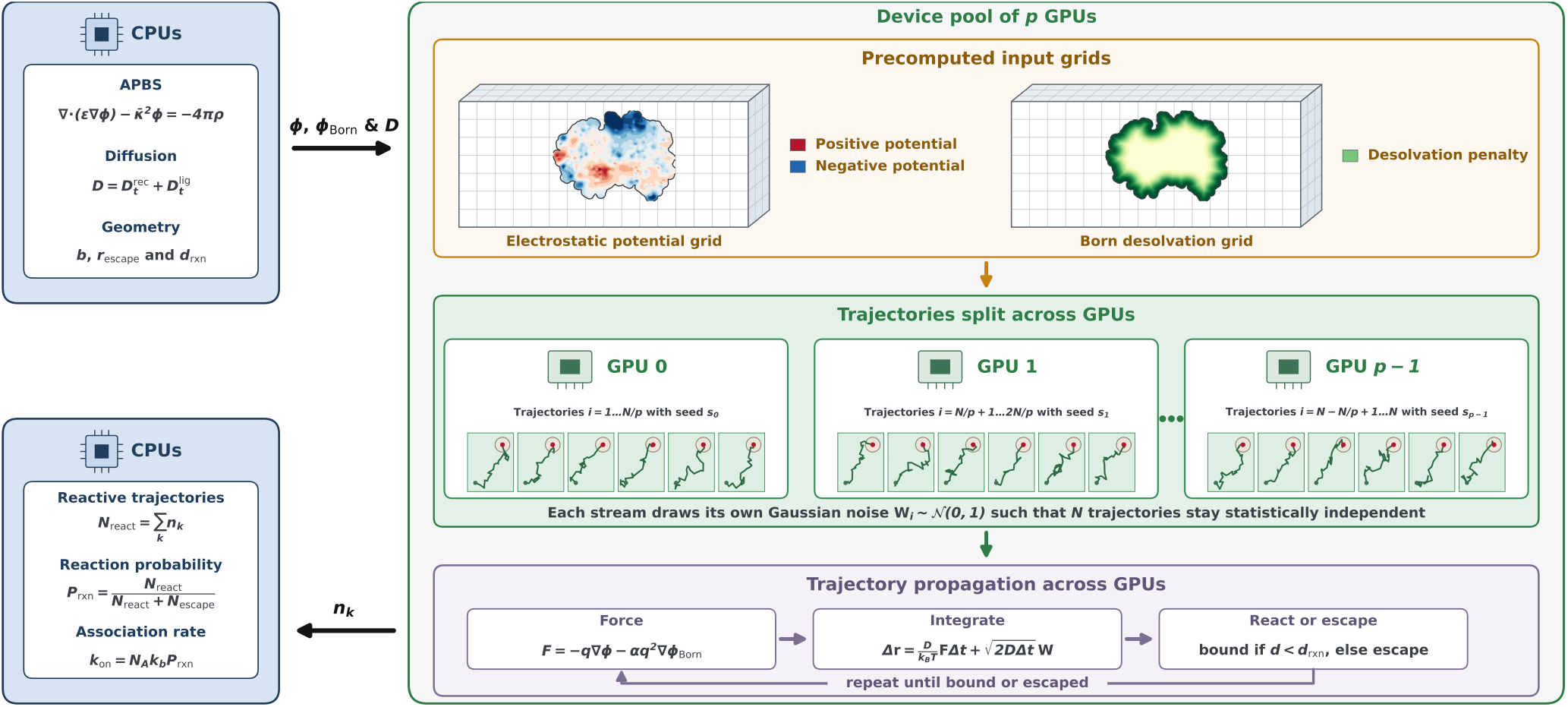
The linearized Poisson-Boltzmann equation ^61^ is solved on the CPUs for the electrostatic potential *ϕ* where *ϵ* is the dielectric coefficient, 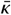 is the modified screening parameter, and *ρ* is the fixed charge density. The relative diffusion coefficient *D* is the sum of the translational coefficients of the two solutes, and the reaction criteria are set by the initiation radius *b*, the escape radius *r*_escape_ and the reaction distance *d*_rxn_. The resulting electrostatic and Born desolvation grids are transferred to a pool of *p* GPUs. *N* trajectories are divided among the *p* devices and advanced with Gaussian increments **W**_*i*_ of zero mean and unit variance, generated independently on each device from its own seed *s*_*k*_. Each trajectory is propagated by the Ermak-McCammon step, where *α* is the desolvation scaling coefficient, until it binds or escapes. The counts *n*_*k*_ are returned to the CPUs and pooled into the reactive and escaped counts *N*_react_ and *N*_escape_, which define the reaction probability *P*_rxn_, scaled by *k*_*b*_ to give *k*_on_.

## RESULTS AND DISCUSSION

*PySTARC* is benchmarked for the trypsin-benzamidine complex on NVIDIA A100 GPUs ^79^, with the number of trajectories *N* varying from 10^4^ to 10^7^ over 1, 2, 4, 6, 8, 12, and 16 GPUs (Figure 3a). Each GPU is initialized independently. The receptor and ligand are centered at the coordinate origin, the b-surface and escape radii are computed from the molecular geometry, the precomputed electrostatic and desolvation grids are loaded into device memory, and the relative diffusion tensor is assembled. At small *N*, each device receives too few trajectories to reach full occupancy, and the propagation is limited by launch and synchronization rather than by the work itself. As a consequence, additional GPUs yield negligible speedup. As *N* increases, the devices saturate, the propagation time scales with the trajectories assigned to each device, and the speedup approaches the device count. At *N* = 10^7^, the wall time falls from 8.98 h on one GPU to 1.04 h on 16 GPUs. The overhead comprises the per-shard setup and the pooling of the per-device outputs. It parallelizes less efficiently than the propagation, and its share rises from 12% for 1 GPU to 37% for 16 GPUs while the overhead time itself decreases. A parallel efficiency of 54% is retained at 16 GPUs (Figure 3a). Since the benchmark simulations differ only in the GPU count and the random seed, any systematic deviation in rate constants would indicate an error in the parallel decomposition. To detect any such deviation, each rate constant is normalized to the single-GPU run at the same *N*. At *N* = 10^4^, normalized rate constants lie between 0.88 and 1.26. The deviation falls below 7% at *N* = 10^5^ and below 1% at *N* = 10^7^. The spread of the normalized rate constants narrows as *N*^−0.511^ against the Monte Carlo limit of *N*^−1/2^, and no bias remains at any GPU count (Figure 3b). For the same trypsin-benzamidine complex, *PySTARC* completes 10^7^ trajectories in 2.65 h on a single GPU node with four A100 GPUs, against 9.9 h for BrownDye2 on the 64 cores of a CPU node, with both engines reproducing the experimental *k*_on_ within the same order of magnitude under their own reaction criterion. A single GPU node therefore replaces approximately 3.7 CPU nodes for this calculation.

**Figure 3:**
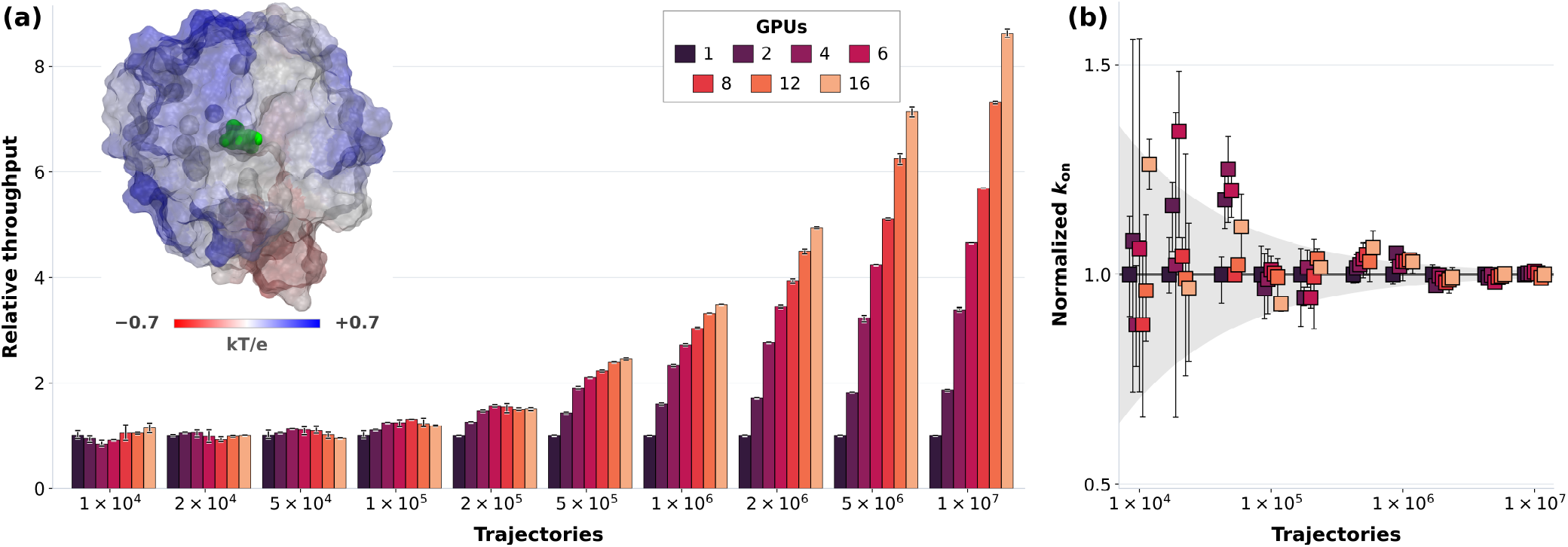
(a) *PySTARC* GPU scaling for the trypsin-benzamidine complex as speedup over a single GPU. The electrostatic potential of the trypsin protein is mapped onto its molecular surface, with the benzamidine ligand in green. (b) The association rate constants obtained from benchmark simulations are normalized to the corresponding single GPU rates. Since only the GPU count and the random seeds differ, agreement with unity validates the parallel decomposition rather than the model. The shaded area is ±1.96 *σ*, with *σ* ∝ *N* ^−0.511^ against the Monte Carlo limit of *N* ^−1/2^.

*PySTARC* is evaluated on complexes that span the physical factors governing bimolecular association, where the association rates span six orders of magnitude, the structural complexity grows from an analytically solvable pair of charged spheres through small-molecule protein-ligand complexes to folded protein-protein interfaces, and the binding sites range from open interfaces to deeply buried selective pockets whose small reaction probabilities make convergence challenging. These complexes span therapeutic areas where association kinetics govern efficacy, including anticoagulation for thrombin and thrombomodulin ^80–82^, chaperone inhibition for heat shock protein 90 (HSP90) ^83–85^, mitotic checkpoint inactivation for threonine tyrosine kinase (TTK) ^86–88^, cytokine suppression for p38 mitogen-activated protein (MAP) kinase ^89–91^, intraocular pressure reduction for carbonic anhydrase ^92–94^, serine protease inhibition for trypsin ^54,95,96^, and targeted cancer therapy for barnase-barstar complex ^97,98^ (Figure 4a and Supplementary Figure S1).

**Figure 4:**
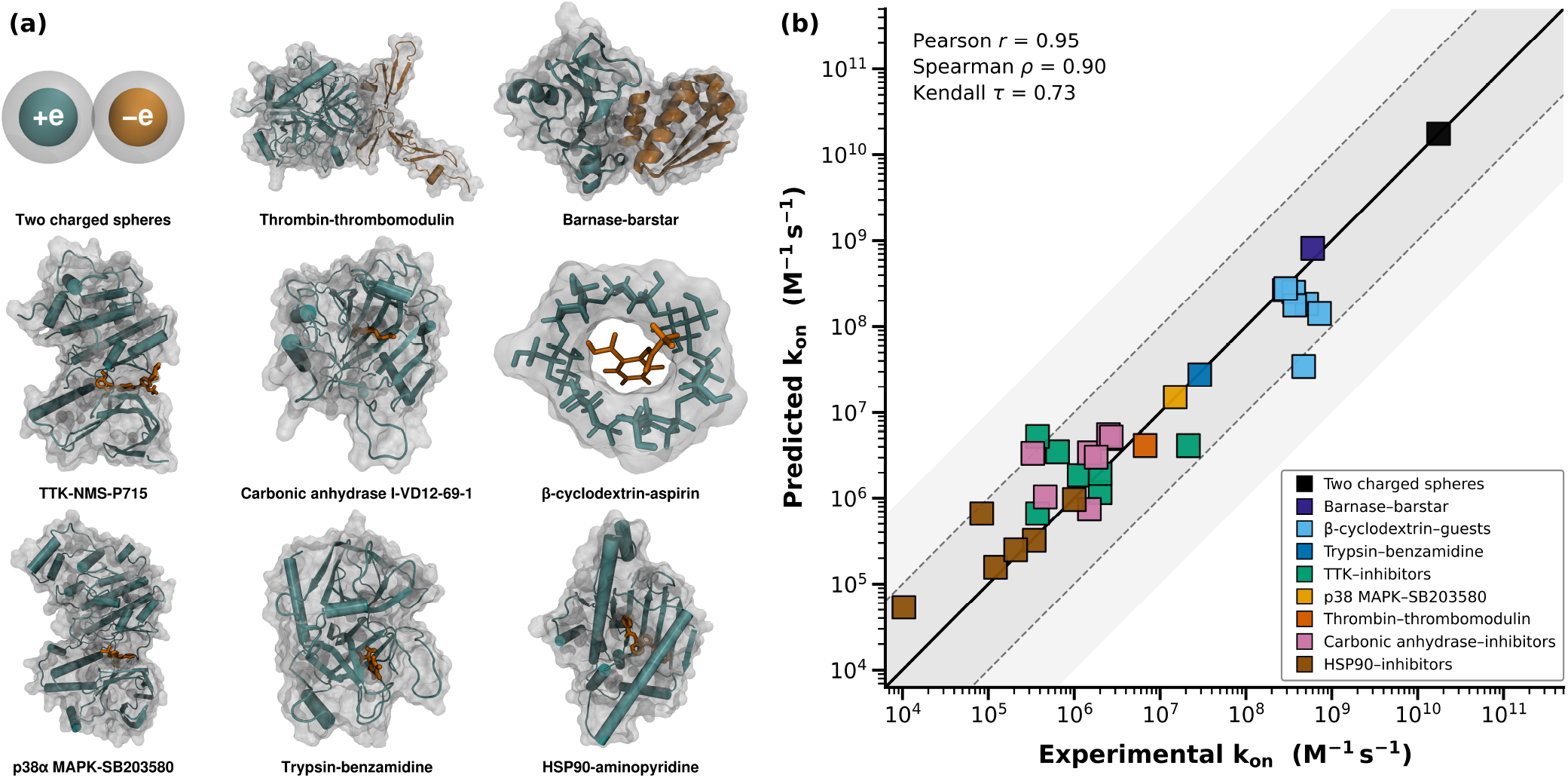
(a) Representative structures of the protein-ligand and protein-protein complexes in increasing order of *k*_on_ rates. (b) Predicted and experimental association rate constants *k*_on_ for the two charged spheres ^44,99^, barnase-barstar ^47,100–102^, *β*-cyclodextrin-guest ^103–108^, trypsin-benzamidine ^109^, threonine tyrosine kinase (TTK)-inhibitor ^110^, p38 mitogen-activated protein kinase (p38 MAPK)-SB203580 ^111^, thrombin-thrombomodulin ^112^, carbonic anhydrase-inhibitor ^113^, and heat shock protein 90 (HSP90)-inhibitor ^28,114,115^ complexes. The shaded regions indicate deviations within one (dark grey) and two (light grey) orders of magnitude. The predicted and experimental rates correlate with a Pearson coefficient of 0.95 (in log_10_ space), a Spearman coefficient of 0.90, and a Kendall coefficient of 0.73.

A BD simulation requires the reaction criterion, the launch surface, and the solvent model to be set before the run. Given the structure of the bound complex, the criterion is built from polar, nonpolar, or heavy-atom contacts between the solutes within a pre-defined search radius. In cases where no bound pose exists, or a defined chemistry governs binding events, the contact pairs are specified directly. Trajectories are launched from the *b*-surface, whose radius exceeds the combined maximum radii of the two solutes so that the intermolecular potential is isotropic at this separation. The solvent is treated implicitly as a dielectric continuum, and the Debye length that screens the electrostatics is determined by the ionic strength of the experimental buffer. Simulation parameters for all complexes, comprising the *b*-surface radii, trajectory counts, grid dimensions, timesteps and reaction criteria, are provided in Supplementary Table S2.

The two charged spheres serve as the analytical reference, since their rate is known in closed form and is recovered to within 0.6% ^99^. Association in the thrombin-thrombomodulin, trypsin-benzamidine, barnase-barstar, and carbonic anhydrase complexes is electrostatically steered, whereas the largely neutral HSP90 and TTK inhibitors, the p38 MAP kinase inhibitor, and *β*-cyclodextrin guest molecules encounter their targets near the diffusion limit. Across all complexes, the predicted and experimental association rates correlate with a Pearson coefficient of 0.95 (in log_10_ space), a Spearman coefficient of 0.90, a Kendall coefficient of 0.73, and 30 of the 33 complexes fall within an order of magnitude of the experimental rates (Figure 4b and Supplementary Table S3). The predicted association rates are distributed evenly above and below the experimental measurements with a median ratio of 1.01 and show no systematic deviation. Within a receptor class, the ranking is less reliable as the diffusional encounter is nearly identical for each ligand ^47^. The remaining differences arise from short-range desolvation and induced fit ^116^, which lie beyond the resolution of a rigid body continuum model ^58,117^. Association rate differences between distinct receptors arise from the electrostatics and the geometry of the encounter and are therefore reproduced across the full range.

The accuracy of the association rate constants estimated by rigid-body Brownian dynamics is limited by a series of approximations. BD simulations compute the rate of transient complex formation at the diffusion limit imposed by an absorbing reaction criterion ^44,46,118^. Association is diffusion-controlled only when rearrangement of the transient complex is fast relative to its dissociation ^47,119^. The timescale of internal motion relative to the encounter lifetime governs the regime, not a deeply buried binding site ^120–122^. The internal degrees of freedom of the macromolecules are greatly reduced, commonly to rigid body translation and rotation ^58,59,123^. The regions requiring finer treatment must be chosen in advance, with the available resolution specified by the software ^59,60,124,125^. Under a continuum solvent of nonuniform permittivity, the inter-molecular forces are many-body rather than pairwise ^126–128^. The electrostatic desolvation term is therefore an approximation of an approximation, i.e., a truncated single-sphere expression summed pair-wise and scaled by a coefficient set to unity throughout ^51,58,59,116,129^. Hydrophobic forces are a major determinant of association rates but are computed only approximately as the product of an empirical surface tension and the buried surface area ^59,129–131^. Hydrodynamic interactions are approximated at several levels of accuracy by the RPY bead models ^56,132,133^. Their influence on the rate and the cost of the mobility tensor both increase with the number of beads ^134–137^. Reaction criteria, including the per-pair contact distances, have been treated as adjustable parameters tuned against measured rates in known complexes and are therefore uncertain to transfer to novel complexes ^51,57,75,138^. Sampling limitations remain for complex macromolecules, though far less severe than in atomistic MD simulations, and are reduced further by enhanced sampling approaches ^34,35,139^. Established multiscale approaches replace the geometric criterion with MD simulations, partitioning the space around the receptor and propagating only the innermost region atomistically ^39,117,140,141^.

A rigid body description is inadequate for solutes with intrinsically disordered regions or flexible loops. The current framework allows either solute to be propagated as a flexible chain of one bead per residue, built from its sequence under the COFFDROP coarse-grained force field for disordered proteins ^142,143^. Neighboring beads are held by harmonic bonds, and the angle, dihedral, and nonbonded interactions are described by the COFFDROP potentials. The chain is evolved using a multiple-timestep Ermak-McCammon integrator, where several short internal steps resolve the fast bonded motion within each longer step that propagates the translation and rotation of the chain ^56^. The overall diffusion tensor of the chain is obtained from an RPY treatment of the beads, while the internal motion neglects hydrodynamic coupling between them ^132,133^. The chain is launched from the *b*-surface and diffuses against the rigid target until the reaction criterion is satisfied, and the trajectories are propagated independently across CPU cores. The barstar protein, represented as a flexible chain, diffuses against a rigid barnase and reproduces the experimental association rate constant to within a factor (Figure 4b and Supplementary Table S3) ^134^.

## CONCLUSION

The association rate can therefore be computed with the solutes held rigid or represented as flexible chains, a choice rather than a constraint ^124^. Each simulation saves a structural record of the encounters, such as first-hitting point distribution at the binding criterion ^76,77^, distance and relative orientation of the two solutes at closest approach, and the radial and angular distributions of the encounters. Machine learning models of binding kinetics require such structure and rate pairs, far scarcer in experiment than affinity data ^70,71,73^. Run across a compound library ^144^, *PySTARC* computes the rate and encounter structure for every complex, and structure-based prediction of association kinetics becomes a question of hardware rather than of measurement.

## DATA AND SOFTWARE AVAILABILITY

The *PySTARC* software is available at https://github.com/anandojha/PySTARC. Tutorials in the examples directory of the *PySTARC* repository demonstrate how to prepare and implement *PySTARC* to estimate the association rate constant for complexes of interest. All data supporting this study, including the *PySTARC* configuration files, the PDB structures and PQR files of solutes, and simulation output files such as the association rate estimates, first passage-time distributions, and bound poses for the 33 complexes, are available at https://doi.org/10.5281/zenodo.21894024.

## SUPPORTING INFORMATION

The supporting information file contains the derivation of the bimolecular association rate constant, BD simulation parameters, and predicted and experimental *k*_on_ rates for the complexes.

## ACKNOWLEDGMENTS

The authors dedicate this manuscript to J. Andrew McCammon ^145^, whose pioneering work on the Brownian dynamics of diffusional association made this study possible. Brownian dynamics simulations were performed on the Rusty cluster at the Flatiron Institute, a division of the Simons Foundation. The authors thank Kanika Dhiman for insightful discussions.

