## Supplementary File for "*PySTARC*: GPU-accelerated Brownian dynamics for bimolecular association rate constants"

**Table S1:** Association rate estimation with *PySTARC*

**NAM framework.** The bimolecular association rate constant in the Northrup-Allison-McCammon (NAM) framework is $k_{\text{on}} = N_A \, k_b \, P_{\text{rxn}}$, where $k_b$ is the diffusion limited encounter rate, $P_{\text{rxn}}$ is the reaction probability, and $N_A$ is the Avogadro constant.

**Diffusion.** The Stokes-Einstein relation $D = \mu \, k_B T$, with mobility $\mu = 1/\gamma$ for drag coefficient $\gamma$, and the Stokes drag coefficients $\gamma_t = 6\pi\eta a$ and $\gamma_r = 8\pi\eta a^3$, gives $D_t = \frac{k_B T}{6\pi\eta a}$ and $D_r = \frac{k_B T}{8\pi\eta a^3}$, where $D_t$ and $D_r$ are respectively the translational and rotational diffusion constants, $\eta$ is the solvent viscosity, and $a$ is the hydrodynamic radius. The two solutes diffuse relative to one another with $D = D_t^{(1)} + D_t^{(2)}$, reduced at short separation by the Rotne-Prager-Yamakawa hydrodynamic coupling.

**Electrostatics.** The electrostatic potential $\phi$ solves the linearized Poisson-Boltzmann equation $\nabla \cdot (\varepsilon \nabla \phi) - \bar{\kappa}^2 \phi = -\frac{\rho}{\varepsilon_0}$, where $\rho$ is the fixed charge density, $\varepsilon$ the dielectric, $\varepsilon_0$ the vacuum permittivity, and $\bar{\kappa}^2$ the screening factor set by the ionic strength, which fixes the Debye length $\lambda$.

**Forces.** The total force $\mathbf{F}$ on an atom of charge $q_i$ is $\mathbf{F} = \mathbf{F}_{\text{elec}} + \mathbf{F}_{\text{Born}} = -q_i \, \nabla \phi - \frac{q_i^2}{4\pi} \, \nabla \phi_{\text{Born}}$, where $\mathbf{F}_{\text{elec}}$ is the electrostatic force, $\mathbf{F}_{\text{Born}}$ is the desolvation force, and $\phi_{\text{Born}}$ is the Born desolvation potential.

**Propagation.** With $D$ from the diffusion step and $\mathbf{F}$ from the force step, the overdamped Ermak-McCammon step advances the relative position by $\mathbf{r}(t + \Delta t) = \mathbf{r}(t) + \frac{D}{k_B T} \mathbf{F} \, \Delta t + \sqrt{2D \, \Delta t} \; \mathbf{W}$, where $\Delta t$ is the adaptive time step and $\mathbf{W}$ is a vector of independent standard normals. The orientation advances the same way with $D_r$, and repeating the step generates one trajectory.

**Reaction criterion.** A trajectory is recorded as reacted when at least $n_{\text{needed}}$ contact pairs are counted within one BD step. Pair $k$ is counted when $d_k$ lies below its cutoff $d_k^{\text{cut}}$ at the end of the step or when the substep crossing test below detects a crossing during the step. With $c_k = 1$ for a counted pair and $c_k = 0$ otherwise, the criterion reads $\sum_k c_k \geq n_{\text{needed}}$. Each pair is tested independently, such that the correlation between pairs on the same rigid solute is neglected.

**Substep crossing.** The reaction criterion is evaluated only at step ends, so a contact pair that starts and ends a step outside the cutoff is still counted as crossing with probability $P_{\text{cross}} = \exp\left(-\frac{x_0 x_1}{D_{\text{eff}} \, \Delta t}\right)$, where $x_0$ and $x_1$ are the distances by which the pair separation exceeds the cutoff at the start and end of the step, and $D_{\text{eff}} = D_t + D_r^{(1)} L_1^2 + D_r^{(2)} L_2^2$ is the effective diffusion coefficient of the pair separation, with $L_1$ and $L_2$ as the distances from each centre of rotation to the contact atom.

**Reaction probability.** Every trajectory ends as either reacted or escaped, and the reaction probability is the reacted fraction, $P_{\text{rxn}} = \frac{N_{\text{react}}}{N_{\text{react}} + N_{\text{escape}}}$, where $N_{\text{react}}$ and $N_{\text{escape}}$ are the numbers of reacted and escaped trajectories, respectively.

**Encounter rate.** The encounter rate $k_b$ is the steady-state Smoluchowski rate at which the two solutes reach the $b$-surface of radius $b$ where the trajectories start, $k_b = 4\pi \left[ \int_0^{1/b} \frac{e^{U(1/s)/k_B T}}{D(1/s)} \, ds \right]^{-1}$, where $s = 1/r$ for solute separation $r$, $V(r)$ is the potential of mean force and $D_\parallel$ the position dependent relative diffusion coefficient. The integral is evaluated numerically and reduces to $4\pi D b$ for a point charge.

**Return probability.** A trajectory that leaves the $b$-surface is restarted at the escape radius $r_{\text{escape}}$ rather than discarded, and returns to the $b$-surface with probability $p_{\text{return}} = \frac{k_b(b)}{k_b(r_{\text{escape}})}$, which is $b/q$ for a point charge.

**Analytic limit.** For two point charges, $k_b$ becomes $4\pi D b$, and summing the geometric series of restarts from $q$ gives $k_{\text{on}} = N_A \, \frac{4\pi D b \, P_{\text{rxn}}}{1 - (1 - P_{\text{rxn}}) \, b/q}$, which at $P_{\text{rxn}} = 1$ reduces to the absorbing sphere Smoluchowski rate $k_{\text{on}} = N_A \, 4\pi D b$.

**Wilson interval.** The uncertainty on $P_{\text{rxn}}$ from the finite trajectory count is the Wilson score interval $p^{\pm} = \frac{P_{\text{rxn}} + \frac{z^2}{2n} \pm z \sqrt{\frac{P_{\text{rxn}}(1 - P_{\text{rxn}})}{n} + \frac{z^2}{4n^2}}}{1 + z^2/n}$, clamped to $[0, 1]$, where $p^{\pm}$ are the upper and lower limits on the true reaction probability, $z = 1.96$ at the 95% level and $n = N_{\text{react}} + N_{\text{escape}}$. Since $k_{\text{on}}$ is linear in $P_{\text{rxn}}$, the interval carries through to the rate as $N_A k_b \, p^{\pm}$.

**Figure S1:** Two-dimensional structures of the threonine tyrosine kinase, heat shock protein 90, and carbonic anhydrase inhibitors and the $\beta$-cyclodextrin guest molecules.

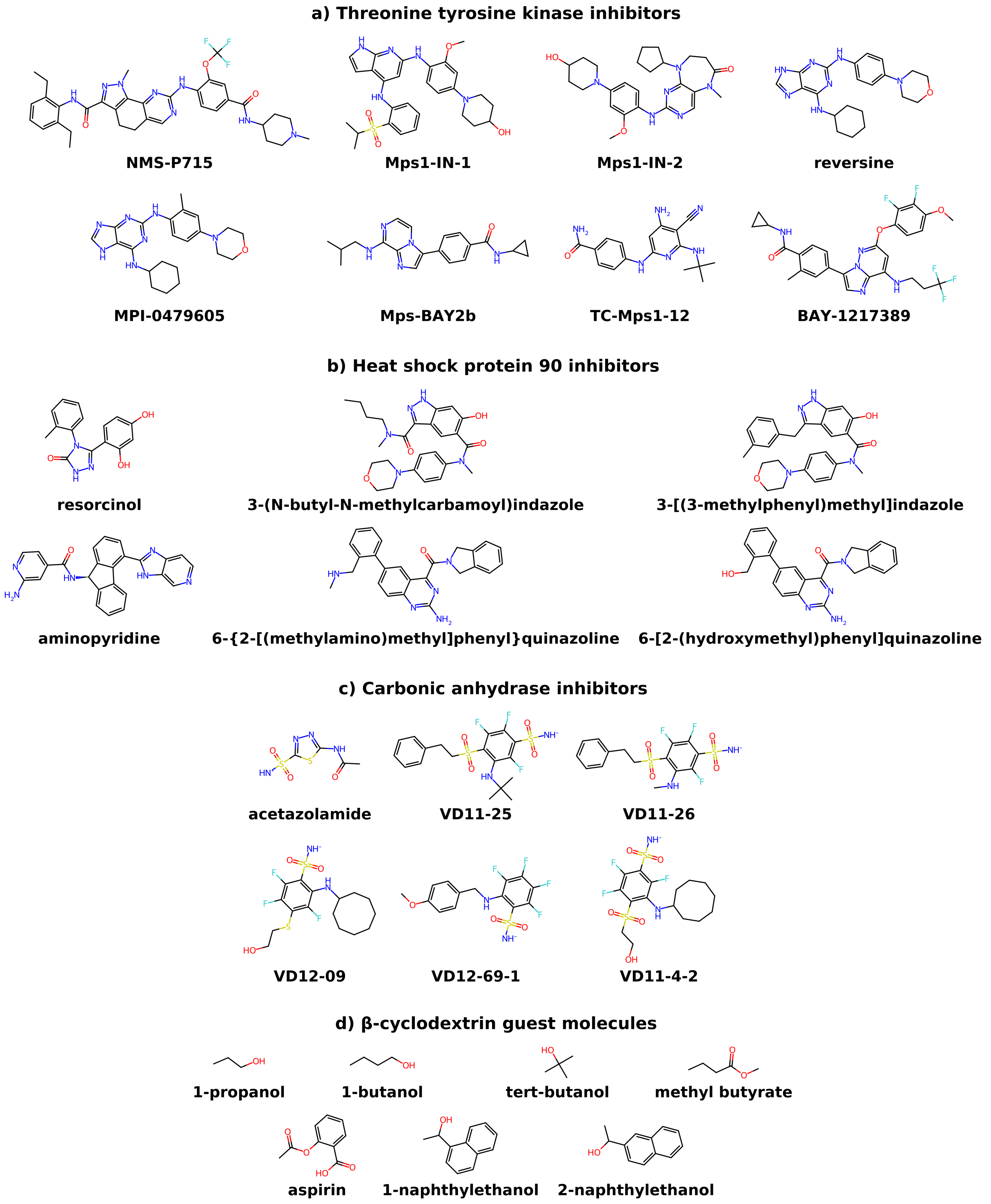

**Table S2:** Simulation parameters for the charged spheres, protein-ligand, and protein-protein Brownian dynamics simulations.

| Complex | $b$-surface (Å) | Trajectories | Maximum steps per trajectory | Timestep (ps) | Hydrodynamics | Solute dielectric | Ionic strength (mM) | Debye (Å) | Temperature (K) | Fine grid (Å) | Spacing (Å) | Reaction criterion |
|---|---|---|---|---|---|---|---|---|---|---|---|---|
| **Analytical validation** | | | | | | | | | | | | |
| Two charged spheres | 10.0 | 1,000,000 | 1,000,000 | 0.2 | False | 78.0 | 150 | 7.83 | 298.15 | Analytical | Analytical | Charged contacts |
| **Protein-ligand complexes** | | | | | | | | | | | | |
| Trypsin-benzamidine | 45.0 | 5,000,000 | 1,000,000 | 0.2 | True | 4.0 | 150 | 7.86 | 298.15 | 96 | 0.375 | Polar contacts |
| p38 MAPK-SB203580 | 60.0 | 5,000,000 | 1,000,000 | 0.2 | True | 4.0 | 150 | 7.86 | 298.15 | 128 | 0.500 | Polar contacts |
| Carbonic anhydrase inhibitors | 60.0 | 20,000,000 | 10,000,000 | 0.2 | True | 4.0 | 100 | 9.62 | 298.15 | 128 | 0.500 | Polar contacts |
| TTK inhibitors | 60.0 | 20,000,000 | 1,000,000 | 0.2 | True | 4.0 | 150 | 7.86 | 298.15 | 144 | 0.562 | Polar contacts |
| HSP90 inhibitors | 55.0 | 20,000,000 | 1,000,000 | 0.2 | True | 4.0 | 150 | 7.86 | 298.15 | 144 | 0.562 | Heavy-atom contacts |
| $\beta$-cyclodextrin guests | 30.0 | 1,000,000 | 1,000,000 | 0.2 | True | 4.0 | 150 | 7.86 | 298.15 | 96 | 0.375 | Heavy-atom contacts |
| **Protein-protein complexes** | | | | | | | | | | | | |
| Thrombin-thrombomodulin | 85.0 | 1,000,000 | 1,000,000 | 1.0 | True | 4.0 | 150 | 7.86 | 298.15 | 192 | 0.750 | Polar contacts |
| Barnase-barstar | 60.0 | 10,000 | 500,000 | 0.2 | True | 4.0 | 150 | 7.86 | 298.15 | 140 | 0.312 | Heavy-atom contacts |

**Table S3:** Predicted and experimental $k_{\text{on}}$ rates for charged spheres, protein-ligand and protein-protein complexes.

| Complex | Experimental $k_{\text{on}}$ ($M^{-1}$ $s^{-1}$) | *PySTARC* $k_{\text{on}}$ ($M^{-1}$ $s^{-1}$) |
|---|---|---|
| **Analytical validation** | | |
| Two charged spheres | $1.75 \times 10^{10}$ [1,2] | $1.76 \times 10^{10}$ |
| **Protein-ligand complexes** | | |
| Trypsin-benzamidine | $2.90 \times 10^{7}$ [3] | $(2.76 \pm 0.06) \times 10^{7}$ |
| p38 mitogen-activated protein kinase (p38 MAPK)-SB203580 | $1.50 \times 10^{7}$ [4] | $(1.52 \pm 0.04) \times 10^{7}$ |
| Carbonic anhydrase (CA)-inhibitor complexes | | |
| CA XIII-acetazolamide | $1.50 \times 10^{6}$ [5] | $(3.42 \pm 0.12) \times 10^{6}$ |
| CA I-VD12-69-1 | $2.70 \times 10^{6}$ [5] | $(5.21 \pm 0.14) \times 10^{6}$ |
| CA XIII-VD12-09 | $3.30 \times 10^{5}$ [5] | $(3.36 \pm 0.11) \times 10^{6}$ |
| CA II-VD11-4-2 | $1.80 \times 10^{6}$ [5] | $(3.04 \pm 0.10) \times 10^{6}$ |
| CA XIII-VD11-26 | $1.50 \times 10^{6}$ [5] | $(7.43 \pm 0.52) \times 10^{5}$ |
| CA XIII-VD11-25 | $4.60 \times 10^{5}$ [5] | $(1.04 \pm 0.06) \times 10^{6}$ |
| CA XIII-VD12-69-1 | $2.50 \times 10^{6}$ [5] | $(5.52 \pm 0.14) \times 10^{6}$ |
| Threonine tyrosine kinase (TTK)-inhibitor complexes | | |
| TTK-reversine | $2.00 \times 10^{6}$ [6] | $(1.16 \pm 0.06) \times 10^{6}$ |
| TTK-NMS-P715 | $6.38 \times 10^{5}$ [6] | $(3.51 \pm 0.10) \times 10^{6}$ |
| TTK-Mps-BAY2b | $2.55 \times 10^{6}$ [6] | $(5.01 \pm 0.13) \times 10^{6}$ |
| TTK-Mps1-IN-1 | $3.73 \times 10^{5}$ [6] | $(6.62 \pm 0.46) \times 10^{5}$ |
| TTK-MPI-0479605 | $2.00 \times 10^{6}$ [6] | $(1.93 \pm 0.08) \times 10^{6}$ |
| TTK-TC-Mps1-12 | $2.14 \times 10^{7}$ [6] | $(4.14 \pm 0.12) \times 10^{6}$ |
| TTK-BAY-1217389 | $3.73 \times 10^{5}$ [6] | $(5.26 \pm 0.13) \times 10^{6}$ |
| TTK-Mps1-IN-2 | $1.14 \times 10^{6}$ [6] | $(1.86 \pm 0.08) \times 10^{6}$ |
| Heat shock protein 90 (HSP90)-inhibitor complexes | | |
| HSP90-resorcinol | $1.00 \times 10^{6}$ [7] | $(9.65 \pm 0.59) \times 10^{5}$ |
| HSP90-3-(N-butyl-N-methylcarbamoyl)indazole | $3.43 \times 10^{5}$ [8] | $(3.21 \pm 0.32) \times 10^{5}$ |
| HSP90-3-[(3-methylphenyl)methyl]indazole | $8.38 \times 10^{4}$ [9] | $(6.60 \pm 0.45) \times 10^{5}$ |
| HSP90-aminopyridine | $1.04 \times 10^{4}$ [8] | $(5.36 \pm 1.29) \times 10^{4}$ |
| HSP90-6-{2-[(methylamino)methyl]phenyl}quinazoline | $1.21 \times 10^{5}$ [8] | $(1.57 \pm 0.22) \times 10^{5}$ |
| HSP90-6-[2-(hydroxymethyl)phenyl]quinazoline | $2.08 \times 10^{5}$ [8] | $(2.50 \pm 0.28) \times 10^{5}$ |
| $\beta$-cyclodextrin (BCD)-guest complexes | | |
| BCD-1-butanol | $2.80 \times 10^{8}$ [10] | $(2.67 \pm 0.04) \times 10^{8}$ |
| BCD-1-propanol | $5.10 \times 10^{8}$ [11] | $(1.84 \pm 0.03) \times 10^{8}$ |
| BCD-tert-butanol | $3.60 \times 10^{8}$ [12] | $(2.53 \pm 0.04) \times 10^{8}$ |
| BCD-methyl butyrate | $3.70 \times 10^{8}$ [13] | $(1.77 \pm 0.03) \times 10^{8}$ |
| BCD-aspirin | $7.20 \times 10^{8}$ [14] | $(1.42 \pm 0.03) \times 10^{8}$ |
| BCD-1-naphthylethanol | $4.70 \times 10^{8}$ [15] | $(3.46 \pm 0.14) \times 10^{7}$ |
| BCD-2-naphthylethanol | $2.90 \times 10^{8}$ [15] | $(2.79 \pm 0.04) \times 10^{8}$ |
| **Protein-protein complexes** | | |
| Thrombin-thrombomodulin | $6.70 \times 10^{6}$ [16] | $(4.14 \pm 0.41) \times 10^{6}$ |
| Barnase-barstar | $6.00 \times 10^{8}$ [17–19] | $(8.14 \pm 0.35) \times 10^{8}$ |
